# Cholesterol depletion by MβCD enhances membrane tension, its heterogeneity and affects cellular integrity

**DOI:** 10.1101/264036

**Authors:** Arikta Biswas, Purba Kashyap, Sanchari Datta, Titas Sengupta, Bidisha Sinha

**Affiliations:** Dept. of Biological Sciences, Indian Institute of Science Education and Research (IISER) Kolkata, Mohanpur - 741246, West Bengal, India

## Abstract

Cholesterol depletion in cells by MβCD remodels the plasma membrane’s mechanics and its interactions with the underlying cytoskeleton. Decoupling the two effects and studying various alterations to the membrane’s mechanical parameters is important for understanding cholesterol’s role in cellular response to stress. By mapping membrane height fluctuations in single cells, we report that MβCD treatment reduces temporal fluctuations and flattens out the membrane – but does not supress activity-driven fluctuations. We find that membrane tension increase contributes most to the altered fluctuations, among the multiple mechanical parameters computed. Maps also reveal an enhanced long-range heterogeneity within single cells, both in amplitude of fluctuations and membrane tension on cholesterol depletion. To check if this alters the tenacity of membrane to mechanical stress we use hypo-osmotic shock. We find that on MβCD treatment, cells are more prone to rupture than control cells, and this is not hindered by actomyosin perturbations. We report increased rupture sizes on cholesterol depletion and argue that, together, this indicates decreased lysis and line tension. Therefore, we show that cholesterol depletion directly affects cell membranes not only by enhancing membrane-cytoskeleton interactions, but also by increasing membrane tension while reducing lysis tension – hence making cells prone to rupture.

## INTRODUCTION

Cholesterol is one of the key components of cell membranes in mammalian cells ^1,2^ and implicated in several cellular functions ^3–8^ including the formation of membrane structures essential for cellular integrity ^9–12^ during stress. Although for some cell types, cholesterol-sensitive structures like caveolae are important ^9,13,14^ for tension regulation during stress, red blood cells (RBCs) -devoid of caveolae- are known to rupture solely by cholesterol depletion^15^. Cholesterol is thus a critical factor in cell membrane tension regulation ^16^ since it can impact the different physical mechanisms used for membrane homeostasis ^17^. In model membranes, cholesterol content not only alters the basic mechanical parameters like bending rigidity ^18^ and elastic modulus ^19^ but also the resistance to rupturing on stress (increasing the line tension ^20^). We therefore ask how cholesterol depletion in cells affects the membrane topology and dynamics; membrane tension and interaction with cytoskeleton; and cellular integrity on stress. Cholesterol is depleted by methyl-beta-cyclodextrin (MβCD) which encapsulates hydrophobic entities of the plasma membrane in its inner hydrophobic cavity ^21^ and extracts cholesterol from the outer leaflet continuously ^22^. Besides its use in cell biology research for cholesterol extraction ^22^, it is also proposed as a drug carrier in anticancer therapies ^23^. While its general effect on tension regulation is not well understood, its impact on membrane mechanics has been extensively studied by micropipette-aspiration and tether force measurements.

Micropipette aspiration studies report increased membrane stiffening on cholesterol depletion but show that the integrity of F-actin is essential for this stiffening ^3^. While tether force measurements show that removal or addition of cholesterol to plasma membranes of cells alters the apparent membrane tension, bending modulus and effective viscosity, increase in membrane-cytoskeleton adhesion is implicated in increasing the tether force ^7,24^. Others techniques have demonstrated that although cholesterol enrichment reduces the membrane-cytoskeleton adhesion, it does not change the global cell stiffness ^7^, probably due to the alteration in the “deep” cytoskeleton rheology ^25^. While in model membranes tether force measurements can lead to a clear understanding of the effect of cholesterol on membrane mechanics, there is no such clarity for plasma membranes in live cells – especially due to the contribution of membrane-cytoskeleton adhesion on the measured apparent membrane tension.

To circumvent this, we use interference reflection microscopy (IRM) to map spatio-temporal membrane fluctuations in live cells and study the effect of MβCD. IRM provides direct information about the membrane topology ^26^, its spatial parameters (correlation lengths) as well as about the dynamics (correlation timescales, power spectral density, standard deviation (SD) of fluctuations and spatial heterogeneity of fluctuations) ^27^. Further comparison with theoretical models help extract the viscoelastic parameters like effective viscosity, membrane tension and confinement parameter. Thus, a detailed picture of the membrane can be accessed by IRM allowing separate evaluations of changes in membrane tension and that in the membrane-cytoskeletal adhesions.

Besides altering the membrane tension, cholesterol content can change the lysis tension in model membranes. Does this also happen in cells? The ability to resist lysis is an important property of lipid membranes in cells ^28^. However, stresses generated or received by organs (flow of fluid on endothelial cells, flow of RBCs, continuous stretching and relaxation of muscles, etc.) can generate physiological ruptures ^29^. Cells rupture when a critical tension (lysis tension) is overcome and the area strain on lipids crosses a threshold. Theoretical studies indicate that while pores of sizes below a critical radius rapidly reseal by line tension, larger pores make the membrane unstable ^20,28^. Thus, pores which are responsible for rupturing reach a critical radius when the critical tension is attained. Membrane rupture has been studied either by electroporation ^20,28^ or photoinduced membrane ablation ^30^. Unlike these techniques that apply local stresses at specific locations on membranes, physiological stresses are global stresses. To understand how cholesterol depletion alters cell membrane integrity, we administer a global mechanical stress (hypo-osmotic shock) on cells and measure the percentage of cells that rupture (propensity) and the kinetics of the decay of trapped fluorophores from the ruptured cells. The latter helps us estimate the rupture pore diameter. Comparing with known models of the lysis tension, line tension and rupture/pore diameter ^20,28,31^, we draw inferences about the effect of cholesterol depletion on line and lysis tension.

## METHODS

### Cell culture and fixation

HeLa (human, female), CHO-K1 (hamster, female) and C2C12 (mouse) cells are grown in Dulbecco’s Modified Essential Medium (DMEM, Gibco, Life Technologies, USA) supplemented with 10% foetal bovine serum (FBS, Gibco) and 1% Anti-Anti (Gibco) and maintained at 37°C in a humidified atmosphere with 5% CO2.

### Preparation of red blood cells (RBCs)

Human RBCs are prepared freshly before each experiment by pricking the finger of a healthy human donor. The blood (collected in 1.5 mL tubes) is centrifuged at 1000 g at 4°C for 10 mins ^32^. The supernatant (consisting of plasma, white blood cells and platelets) is carefully removed. The pellet containing the RBCs is resuspended in 1X Hank’s Balance Salt Solution (HBSS (+ Calcium Chloride, + Magnesium Chloride), Gibco). 150 µl of this resuspended solution is plated on fibronectin (25 µg/ml) coated coverslips and incubated at 37°C for 3 hrs.

### Staining for cholesterol

HeLa cells, seeded at a concentration of about 20,000 cells/ml (between passages 3 and 17) are deposited on customized glass bottomed dishes and all experiments are performed after 16 hrs of seeding. For staining cholesterol, cells are fixed with 4% paraformaldehyde (Sigma, USA) for 15 mins, washed thoroughly with 1X phosphate buffer saline (PBS, Sigma) and then incubated in 0.1 M glycine (Sigma) for 5 mins. They are washed well and then incubated with 0.05 mg/ml Filipin III (Santa Cruz Biotechnology, USA) in the dark for 2 hrs ^33^. Cells are always washed before imaging.

### Pharmacological treatments

Cells are incubated with 5 µM Cytochalasin D (Cyto D, Sigma) for 60 mins to inhibit the polymerization of actin filaments ^34^. To deplete the cells of cellular activity, 10 mM sodium azide (Sigma) and 10 mM 2-deoxy D-glucose (Sigma) are added to cells in M1 Imaging medium (150 mM NaCl (Sigma), 1 mM MgCl2 (Merck, USA) and 20 mM HEPES (Sigma)) and incubated for 60 mins ^35^. For cholesterol depletion, cells are incubated with 10 mM methyl-beta-cyclodextrin (MβCD, Sigma) in FBS free DMEM for 50 mins ^10^. For dual drug treatments, cells in serum-free medium are treated first with Cyto D for 60 mins and then with MβCD for 50 mins without replacing the medium (Cyto D + MβCD). The reverse order of treatments is denoted as MβCD + Cyto D in the study. All the incubations are done at 37 °C.

### Fluctuations based experiments

Cells are imaged in an onstage 37 °C incubator (Tokai Hit, Japan) atop a Nikon Eclipse Ti-E motorized inverted microscope (Nikon, Japan) equipped with adjustable field and aperture diaphragms, 60X Plan Apo (NA 1.22, water immersion), a 1.5X external magnification and an EMCCD (Evolve 512 Delta, Photometrics, USA). For IRM, an additional 100 W mercury arc lamp, an interference filter (546 ± 12 nm) and a 50-50 beam splitter is used as described in ^27^ and time-lapse images are recorded at EM gain 30 and exposure time 50 msecs for 102 secs at 19.91 frames/sec (2048 frames).

### Hypo-osmotic shock induced rupture experiments

Cells (HeLa and RBCs) are incubated with 2.5 µM Calcein AM (Invitrogen) at 37 °C for 30 mins. They are washed well before fresh medium (with/without drugs) is added for further experiments. Epifluorescent images are acquired on the same microscope and camera (used above) with a 10X Plan Apo (NA 0.45, dry) and a 1.5X external. To calculate rupture diameter, image stacks of Calcein AM loaded cells are captured (using the FITC filter set) at 100 ms and 0.5 frames/sec for 5 mins. A mixture of DMEM and deionized water (in the ratio 1:19, 95% hypo-osmotic shock) is then added to the cells and image stacks with the same acquisition settings are captured. For RBCs, the hypo-osmotic shock is 67% and acquisition rate is 2 frames/sec. To calculate rupture propensity, dishes of Calcein AM loaded cells are scanned and multiple fields of these cells are captured in the differential interference contrast (DIC) and epifluorescence modes 15-30 mins after the hypo-osmotic shock administration.

### Calculation of spatio-temporal fluctuations parameters

MATLAB is used to calculate the relative height of basal plasma membrane of the cell from the intensities in each pixel of an IRM image by comparing with images of beads (60 µm diameter polystyrene beads, Bangs Labs.) imaged on the same day as explained in ^27^. Parameters of temporal fluctuations and spatial undulations in the first branch regions (FBRs, these are regions lying in the first branch of the interference pattern and limited to 12 × 12 pixels for consistency of analysis) are then calculated as in ^27^. The parameters of spatial undulations include SD(space) (calculated from standard deviation (SD) of relative heights across 144 pixels in an FBR, averaged over 20 frames) and correlation length, λ (calculated from spatial autocorrelation functions (ACFs) across 350 pixels, averaged over 200 frames). The parameters of temporal fluctuations comprise of mean relative height, SD(time) (mean and SD of relative heights calculated respectively in each pixel over 2048 frames and averaged across 144 pixels), 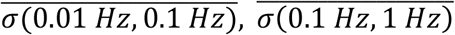 (calculated as root of the total area under the power spectral density (PSD) curve between the mentioned frequencies), exponent (calculated as the linear slope between 0.04-0.4 Hz of the PSD in the log-scale), f (calculated as the ratio between background subtracted PSD of treated to control) and correlation time, τ (calculated from temporal ACFs over 2048 frames, averaged over 4 pixels). The Gaussian-ness of temporal fluctuations is evaluated at each pixel by the Kolmogorov-Smirnov hypothesis testing. Mechanical parameters A, ηeff, γ, κ, µ and σ are computed from fitting the PSDs of FBRs to 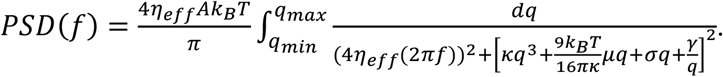. Short-range heterogeneity in cells is measured by SD(SD(time)) (calculated from SD of SD(time) of pixels over 2048 frames and across 144 pixels) and long-range heterogeneity is calculated by percentage of dissimilar FBR pairs (calculated as ratio of number of dissimilar FBR pairs (p-values < 0.001) to total number of FBR pairs; the SD(time) of all 144 pixels in an FBR is statistically compared to those in every possible FBRs in pairs, and p-value from a one-way Analysis of Variance (ANOVA) is calculated). Dissimilar FBR pairs in terms of mean relative height are evaluated from the ratio of the number of FBR pairs with dissimilar mean relative heights (p < 0.001, one-way ANOVA) to the total number of FBR pairs. The log(SD ratio) or log(σ ratio) is computed as the ratio of SD(time) or σ between FBR pairs (bigger value to smaller value) in both dissimilar and similar sets.

### Calculation of rupture propensity

15-30 mins after the application of a hypo-osmotic shock, the total number of cells (Nt) present in each field is counted from the DIC images. The total number of Calcein AM loaded fluorescent cells (Nnr) in the same field is counted from the epifluorescent images and the rupture propensity (RP) in that field is calculated as:

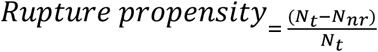

### Calculation of rupture diameter

The lipophilic and non-fluorescent Calcein AM can permeate inside live cells. In the presence of esterases in such cells, the acetoxymethyl group are cleaved trapping the Calcein (not lipophilic and fluorescent in green) molecules ^30,36^. The trapped Calcein moves out of the cell when it ruptures and hence a rupture corresponds to a sudden drop in the Calcein intensity in the cell.

A model based on simple diffusion (Calcein moves out of the cell by diffusion through the rupture site) is used to calculate the rupture diameter ^30^. This model assumes a cell with membrane thickness (l) and a volume (V) to undergo a single point rupture and the rupture pore is assumed to be a cylinder, of length l, radius r and cross-section area A=r^2^. To determine the expected temporal evolution of Calcein concentration (c(t)) in the cell, the expected flux (j) is compared with Fick’s law. Flux, the number of Calcein particles crossing the membrane per unit area per unit time is expressed as 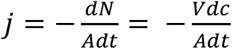. By Fick’s law, we know, 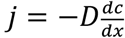, where D is the diffusion constant of Calcein and 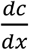 he concentration gradient across the pore. Or, 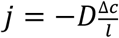 where Δ*c* = *c_out_* − *c_in_* = −*c* assuming Calcein concentration outside to be 0. Therefore, 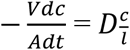 or, 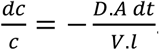. Integration yields, c(t) 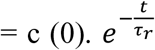 where, 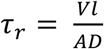. Therefore, the concentration evolution of the flow of Calcein from inside to outside follows an exponential.

A region inside the ruptured cell is selected and its normalized mean intensity (mean intensity of region in each frame is divided by the mean intensity in the first frame) is plotted with time using a program written in MATLAB. Two points are chosen such that one marks the start of this drop and the other marks the trailing end. All data points within this region are fitted to a double exponential, *f*(*t*) = *Ae^−bt^* + *Ce^−dt^*. The damping constant b is used to calculate 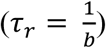 while the other represents the photobleaching in the system, if any. Assuming the radius (R) of a typical HeLa cell is 20 µm (and 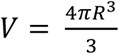 and V = 100 µm^3^ for RBCs), l is 7 mm and D is 330 µm^2^/s, the rupture diameter, *r_D_* (2r) is calculated from: 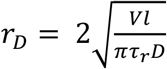.

### Statistical Analysis

For fluctuations-based experiments, calibration with beads and control experiment with cells without any treatment are performed with each set of experiment. At least 10 cells are imaged for each condition and ∼20-40 FBRs analysed for each cell. In most cases, analysis is collated over at least three sets of experiments performed on different days. For comparisons between populations of cells, a one-way ANOVA combined with a Tukey post-hoc test is performed to determine the statistical significance (* denotes p < 0.05, ** denoted p < 0.001) whenever the parameters have similar variances and have Gaussian distributions. A Mann-Whitney U test is done whenever the parameters are not Gaussian (checked if the mean and median values are not similar). The experiments with hypo-osmotic shock are done on at least three days. Rupture diameter is calculated from multiple cells in a single field in each day and rupture propensity is calculated from multiple fields imaged in a day. Values of rupture propensity higher than mean + 2SD are considered as outliers are removed for the box plots. A Mann-Whitney U test is done to determine the statistical significance (* denotes p < 0.05, ** denoted p < 0.001). No specific tests are done to check the normality of the data.

## DATA AVAILABILITY

The datasets generated and analysed during the current study are available from the corresponding author on reasonable request.

## RESULTS

### MβCD treatment decreases temporal fluctuations and flattens out spatial undulations

HeLa cells are depleted of cholesterol by MβCD and stained with Filipin III to check for cholesterol depletion (Fig. 1 A, *left*). As seen in earlier reports ^37^, images show filipin staining at the plasma membrane in control cells which is lost on MβCD treatment, thereby, increasing the contrast of the intracellular vesicles. A quantification of mean Filipin III fluorescence at the membrane shows a ∼40% (N = 30 cells) decrease on cholesterol depletion (Fig. 1 A, *right*). Imaging cells in the IRM mode (Fig. 1 B, *left* and Movie S1) reveals that the topology and dynamics of the basal plasma membrane is significantly altered on MβCD treatment. The changes in the dynamics can be visualized by the color-coded kymographs (Fig. 1 B, *middle*) which show reduced variations after cholesterol depletion. Next, we convert intensity in the images to relative heights ^27^ and quantify various spatio-temporal parameters of the height-fluctuations. The amplitude of the temporal fluctuations (SD(time)) reduces on MβCD treatment. This is evident from SD(time) maps (Figs. 1 B, *right* and S1 A-B) as well as from statistics obtained from ∼1500 first branch regions (FBRs) across ∼70 cells per condition. There is a slight but significant decrease in the amplitude obtained from spatial maps (SD(space)) for the same sets of cells (Fig. S1 C). Together, these data imply that spatio-temporal fluctuations are damped on cholesterol depletion. Though the reduction in power-spectral density (Fig. 2 A) is observed to be more prominent at lower frequencies (∼ 0.01-0.1 Hz) (Fig. 2 A, *inset*), the calculated amplitudes (𝜎̅) at both frequency bands – 0.01-0.1 Hz and 0.1-1 Hz –show a significant reduction on cholesterol depletion (Fig. 2 B). The PSD’s power-law dependence on frequency, captured by the exponent, increases (from -4/3 to -1) on MβCD treatment (Fig. S1 D), implying increased damping ^38,39^.

**Figure 1:**
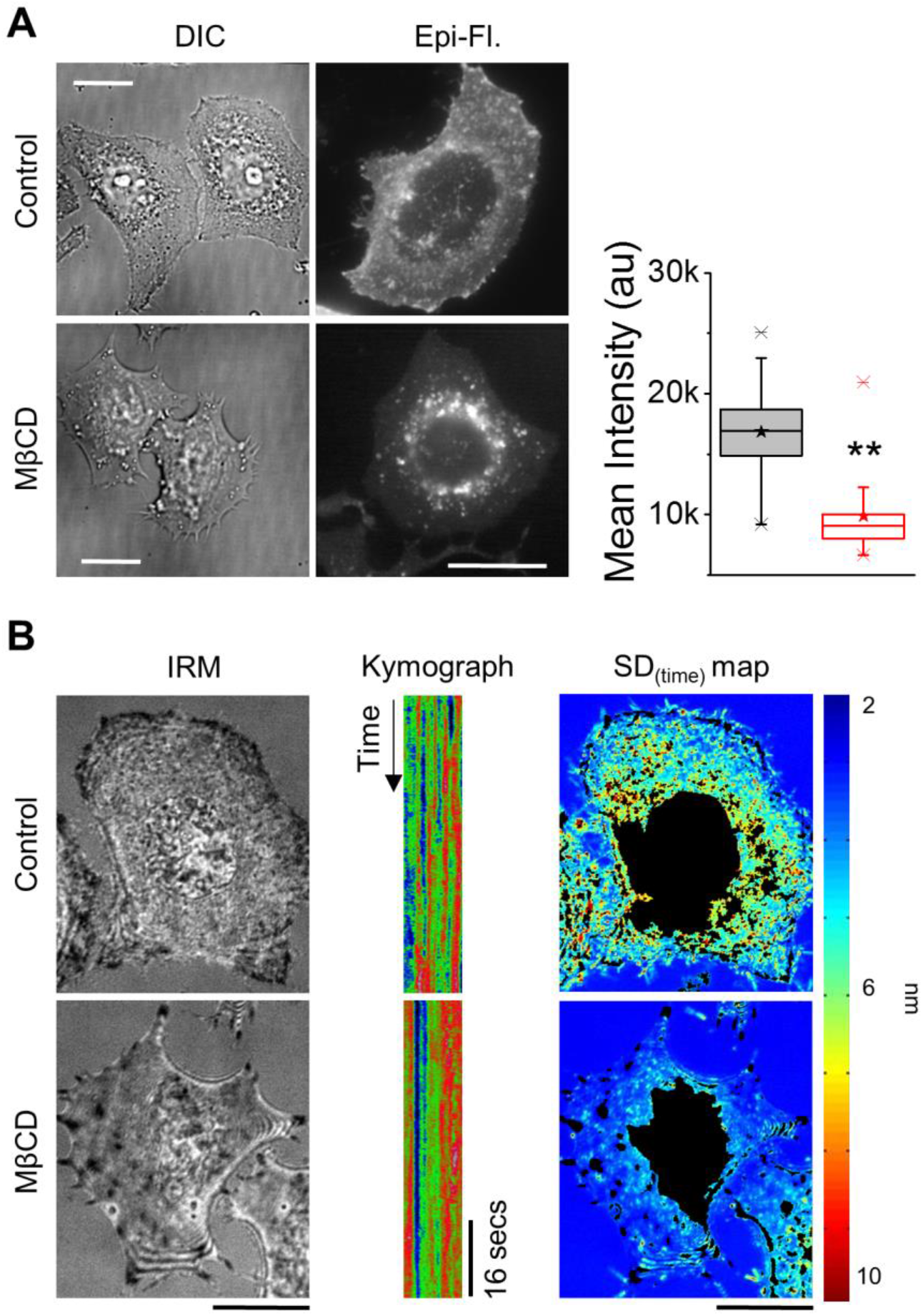
Effect of MβCD mediated cholesterol depletion on membrane topology. A) Representative DIC and epifluorescent images of Filipin III stained control and MβCD treated HeLa cells. Right: Box plot of mean intensity of Filipin III in cells in mentioned conditions (N = 30 cells each). Centre lines of boxes show the medians; stars show the means; boxes limits indicate the 25th and 75th percentiles; and whiskers extend 1.5 times the interquartile range from the 25th and 75th percentiles. ** p value < 0.001, Mann-Whitney U test. B) Representative IRM (left) images, kymographs of 1 µm regions (middle, scale bar: 16 secs) and SD(time) maps of control vs. MβCD treated cells (non FBRs blacked out). Scale bar: 10 µm. See Table S1 for statistics.

**Figure 2:**
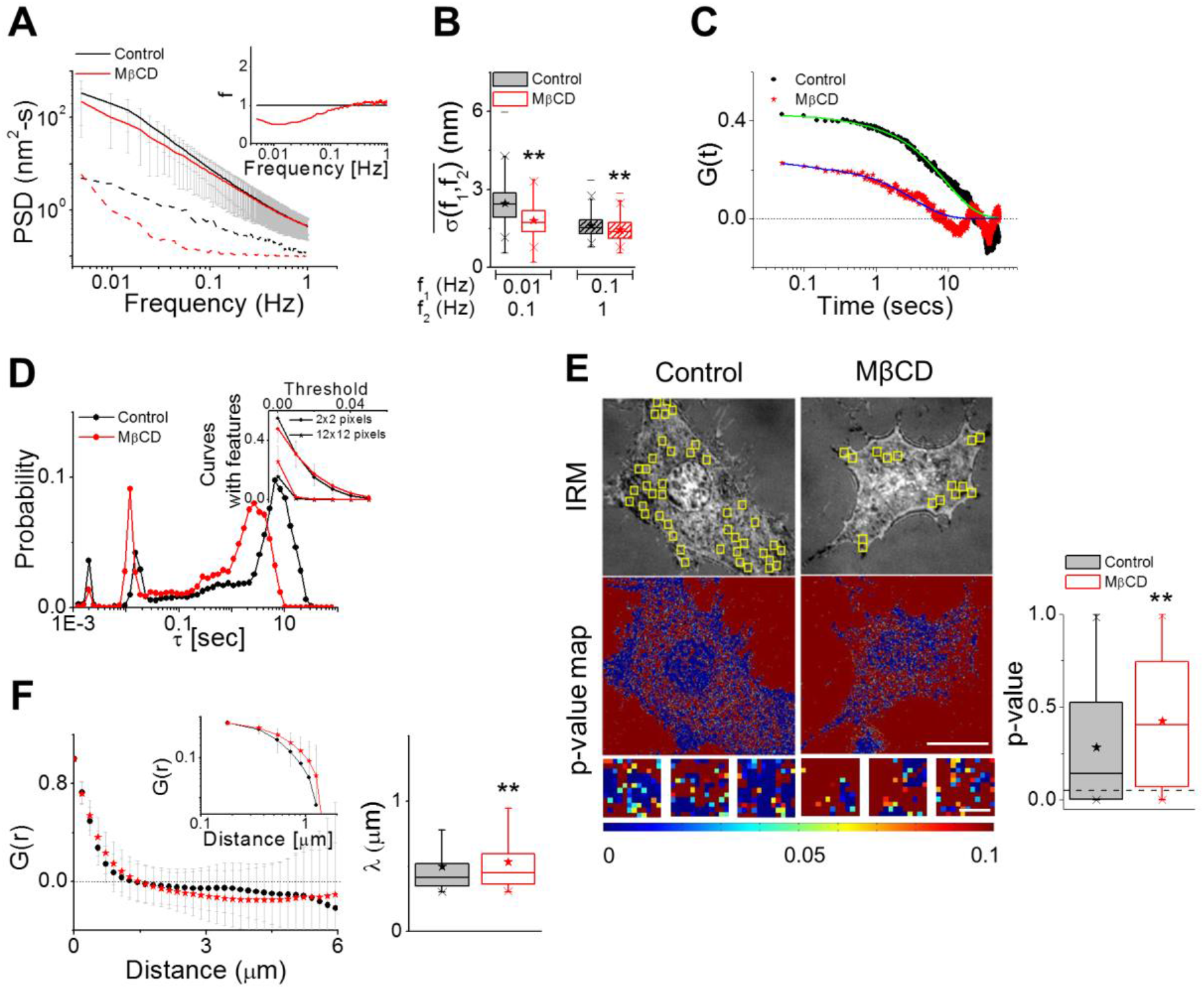
Cholesterol depletion by MβCD reduces temporal fluctuations without abrogating the signatures of activity. A) Averaged PSDs of FBRs in MβCD treated cells and their controls (solid lines) with their backgrounds (dashed lines); inset shows f (ratio of background subtracted PSDs). B) Box plots of 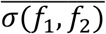 in two different frequency regimes. (A-B) N = 70 cells each, ncontrol = 1688 FBRs, nMβCD = 1474 FBRs. ** p value < 0.001, one-way ANOVA. C) Typical temporal ACFs of single 2 × 2 pixels FBRs in the two conditions. D) Weighted distribution of correlation timescales obtained from temporal ACFs. Inset shows a plot of fraction of curves with features vs. threshold used to detect the features (ncontrol = 2890 fits, nMβCD = 3071 fits, N = 21 cell each). E) FBRs overlaid in yellow on IRM images and their corresponding whole cell and FBR (Scale bar: 1 µm) p-value maps (Kolmogorov-Smirnov hypothesis testing). Right: p-value for FBRs in control vs. cholesterol depleted cells. ncontrol = 53568 pixels, nMβCD. = 42480 pixels. F) Averaged spatial ACFs (and their log-log plots, top inset) for control and cholesterol depleted cells (N = 70 cells, ncontrol = 624 FBRs, n MβCD = 541 FBRs). Right: Correlation lengths. *p value < 0.05, ** p value < 0.001, Mann-Whitney U test.

We next analyse the temporal data to understand how cholesterol depletion affects the contribution of activity-driven processes to the observed fluctuations. We, first, compute the temporal ACFs and find reduced correlation strengths on MβCD treatment (Fig. 2 C). While this is expected, since reduced fluctuations decrease the signal to noise ratio, we also check for signatures of active fluctuations by extracting the correlation timescales (τ). Surprisingly, the distribution of correlation timescales still retains peaks in the 0.2-2 sec range (Fig. 2 D). This range is usually under-represented when ATP-dependent metabolic activity is hampered by ATP depletion ^27^ The other signature of active fluctuations is the presence of “bumps” in ACFs which are known to be affected by ATP depletion ^27,40^ On cholesterol depletion, in this study, we find that similar proportions of ACFs have “bumps” (Fig. 2 D, *inset*). Increasing the region of averaging from 2 × 2 pixels (0.36 µm × 0.36 µm) to 12 × 12 pixels (2.14 µm × 2.14 µm), we find the proportion of ACFs with bumps to decrease –suggesting that localized ATP-dependent processes still affect membrane fluctuations even after cholesterol is depleted. Is the level of “Gaussian-ness” of fluctuations in these cells also retained? We map the p-values of Kolmogorov-Smirnov hypothesis testing to quantify similarity of the temporal fluctuations at each pixel with Gaussian distributions – where higher p-values indicate greater similarity to Gaussian fluctuations (Fig. 2 E). We find that the p-values increase significantly on MβCD treatment. Such increase may result either from the loss of ATP-dependent fluctuations ^27,41^ or may be a result of a reduction in the strength of fluctuations. On analysing data for mitotic cells, where, fluctuation-strength reduces with respect to interphase cells, we find an increase in Gaussian-ness. These cells are expected to retain ATP-dependent activities as also corroborated by existence of correlation timescales at 0.2-2 secs. The level of Gaussian-ness is, thus, determined more by the strength of the fluctuations than by the presence/absence of ATP-dependent fluctuations. Together, this indicates that cholesterol depletion reduces fluctuations without removing activity-driven fluctuations.

To understand the alterations to the spatial membrane topology, we first quantify the changes in relative height. On cholesterol depletion, the average membrane-substrate distance (and its variability) increases significantly (Fig. S1 E). Autocorrelation function of the spatial height profiles (Fig. 2 F) show that the correlation length increases significantly (from 0.49 ± 0.27 µm to 0.53 ± 0.28 µm) on MβCD treatment. Together with decreased SD(space), this implies that cholesterol depletion spatially flattens the membrane. The trends of altered spatio-temporal parameters are checked in MβCD treated- CHO and C2C12 cells (Fig. S1 F-I). As seen in HeLa, the other cell lines show reduced amplitudes of temporal fluctuations. Exponent, mean relative height and correlation length also increase on MβCD treatment in these cell lines (like HeLa).

### Cholesterol depletion by MβCD increases membrane tension and its heterogeneity

We next compute membrane mechanical properties by fitting the PSDs with a theoretical model ^27^ to further characterize the effect of cholesterol depletion. The model predicts the PSD for a confined (confinement, γ) membrane of defined tension (σ), bending rigidity (κ) and shear modulus (µ) in a viscous surrounding (of effective viscosity, ηeff) and acted on by active forces to increase the temperature “A” times ^38,40,42,43^. On fitting, we find that while A decreases, ηeff, γ, µ, and σ increases significantly on MβCD treatment (Fig. 3 A). Since multiple parameters are affected, we next sought to understand which one is the most important in altering the fluctuation amplitude. For this, we compared the SD of the control set with the SD calculated from simulated PSDs. The simulated PSDs are generated from different sets of observed fitting parameters. We chose a whole set of fitting parameters corresponding to a control set and then changed only one parameter at a time to that of a MβCD treated set. This is done for each of the parameters – A to σ. The ratio of the SD of the original control set and the SD calculated from the simulated PSD is calculated and the log of these values are plotted to understand the different contributions. We find that with all other parameters being same to the control, changing σ has the most significant effect in the reduction of SD. Hence, we believe that although multiple parameters are altered on MβCD treatment, alterations in membrane tension govern the change in fluctuations (Fig. S2 A).

**Figure 3:**
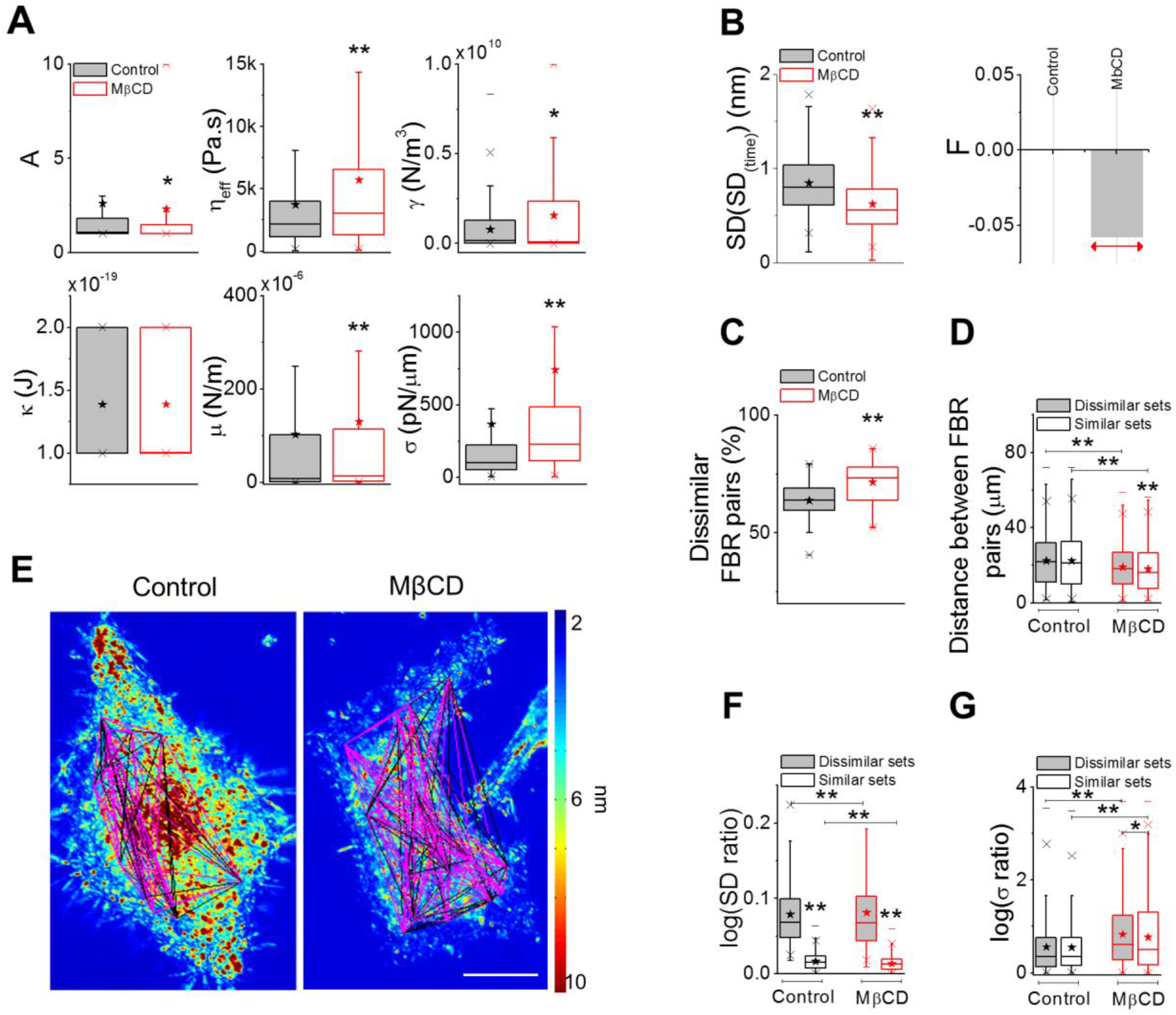
Membrane mechanical parameters and tension heterogeneity in single cells. A) Membrane mechanical parameters A, ηeff, κ, µ, σ and γ obtained from fitting PSDs to theoretical model (ncontrol = 1031 FBRs, nMβCD = 977 FBRs, N = 70 cells each) B) SD(SD(time)) for the two conditions (left) and the Fano factor (right). ncontrol = 1688 FBRs, nMβCD = 1474 FBRs, statistical testing by one-way ANOVA. C) Box plot of the number of dissimilar FBR pairs evaluated by comparing SD(time). Statistical testing by one-way ANOVA. D) Distance between FBR pairs that have dissimilar (grey filled) and similar (no filled) SD values in the two conditions. ncontrol = 15158 dissimilar sets, 8148 similar sets; nMβCD = 12451 dissimilar sets, 4986 similar sets. E) Lines in magenta and black connect FBRs that are dissimilar and similar in SD values respectively. The lines are overlaid on the SD(time) maps and each node represents the centre of an FBR. F) Box plot of log SD ratio of dissimilar and similar sets in control vs. cholesterol treated cells. G) Box plot of log σ ratio of dissimilar and similar sets in the above-mentioned conditions. ncontrol = 15135 dissimilar pairs, 8171 similar pairs; nMβCD = 12713 dissimilar pairs, 4724 similar pairs. N = 70 cells each. * p value < 0.05, ** p value < 0.001, Mann-Whitney U test. Scale bar: 10 µm. See Table S1 for statistics.

We next seek to understand the effect of cholesterol depletion on the spatial heterogeneity and characterize both short (inside an FBR, <2.14 µm) and long-range heterogeneity (distances ranging from 2.14 µm – 54 µm). To compute short-range heterogeneity, we calculate the SD(SD(time)) and find a reduction in this quantity after MβCD treatment (Fig. 3 B, *left*). To normalize out the effect of the reduced mean, we calculate the Fano factor of the fluctuations and find a reduction on cholesterol depletion (Fig. 3 B, *right*). Long-range heterogeneity is calculated by comparing all possible pairs of FBRs in cells and obtaining the p-values (of SD(time)s) to segregate similar (p > 0.001) and dissimilar (p < 0.001) FBR pairs. The percentage of dissimilar FBR pairs increases on cholesterol depletion (Fig. 3 C). To understand what factors, lead to this increased dissimilarity, we calculate the distance between FBR pairs for all similar and all dissimilar sets – and compare the mean values of each set. In control cells, there is no significant difference in the distance between FBR pairs in similar and dissimilar sets. However, the dissimilar set shows significantly higher mean distance than the similar set in the cholesterol depleted cells (Fig. 3 D). We visualize the sets (similar: magenta lines, dissimilar: black lines) in both conditions to understand if there is a correlation between dissimilarity and their location in cells. We find that neither sets are located at a specific region in the cells – e.g. cell periphery, perinuclear regions, etc. (Figs. 3 E and S2 B) in any of the two conditions. We next check if membrane-substrate distances are responsible for the observed increase in long-range heterogeneity. For this, we compute p-values for all FBR pairs based on their mean relative heights and find that although dissimilar pairs exist (∼62%) – this does not change on cholesterol depletion (Fig. S2 C, *left*). For these sets of similar and dissimilar FBR pairs, the distance between pairs do not vary significantly in either of the conditions (Fig. S2 C, *right*).

Again, visualizing the connections show that there is no striking correlation to their underlying mean height profile (Fig. S2 D). Therefore, the increased spatial heterogeneity is neither driven by altered adhesion state nor by specific intracellular localization. It is important to note that such long-range spatial heterogeneity is not increased by ATP depletion or cytoskeletal perturbations, as observed earlier by our group ^27^. To check if the dissimilar and similar sets have different values of SD and σ, we find the SD(time) ratio (calculated between the bigger and smaller value for each pair) in every pair of the two sets in each condition. We find that in control and MβCD treated cells, the SD ratio is smaller for similar pairs than the dissimilar ones (Fig. 3 F). On computing the σ ratio, in the same way, we find that the dissimilar sets in cholesterol depleted cells have a higher tension than the similar sets (Fig. 3 G). This analysis, along with the variability in tension on MβCD treatment (Fig. 3 A) confirms that the increased spatial heterogeneity in fluctuations maps on to the increased heterogeneity in tension. However, the increase in σ ratio on cholesterol depletion observed in both similar and dissimilar sets points out that the tension heterogeneity is amplified independent of the corresponding SD ratios.

There is a decrease in SD(SD(time)) in CHO cells depleted of cholesterol (as in HeLa cells) but there is no significant change in the short-range heterogeneity in cholesterol depleted C2C12 cells (Fig. S2 E, *left*). The long-range heterogeneity does not show any significant change on cholesterol depletion in CHO or C2C12 cells (Fig. S2 E, *right*). But, it is noteworthy that, in both cell lines, MβCD treatment leads to an increase in σ and its variability (Fig. S2 F).

We compare γ values between control and MβCD treated cells to check if our results show increased membrane-cytoskeleton adhesion as reported earlier ^44^. We find that, for HeLa cells, there is a significant increase in γ on MβCD treatment. γ defines the overall confinement - due to membrane-cytoskeleton as well as membrane-substrate interactions. Since cholesterol depletion increases the membrane-substrate distances (Fig. S1 E), we believe that the amplification in γ is due to increased membrane-cytoskeletal adhesion. The increase in damping (higher exponent) and effective viscosity, also, support this inference. In CHO and C2C12 cells, γ is not significantly altered by MβCD treatment. The increase in mean relative height is also larger in these cell lines, which we believe, may cancel out the effect of membrane-cytoskeleton attachment. Though the overall increase in membrane-cytoskeleton adhesion is clear in HeLa cells, the effect is not robust across different days of experiments or across cell lines. While IRM helps us to estimate membrane tension, we acknowledge that we cannot separate out the contribution of confinement by the cytoskeleton from that of damping by the substrate.

Therefore, the most robust effect of MβCD on membrane mechanics in single cells is to increase the membrane tension and its spatial heterogeneity. In the next section, we address how MβCD treatment alters the rupture propensity and affects the lysis tension in cells.

### Hypo-osmotic shock induced membrane rupturing propensity and rupture diameter increases on MβCD mediated cholesterol depletion

We use hypo-osmotic shock to impart global mechanical stress (Movies S2 and S3) on cell membranes and assess its propensity to rupture. We load HeLa cells with Calcein AM and analyse the cells before and at least 15 mins after hypo-osmotic shock. Cells with ruptured membrane lose the internal Calcein AM and are hence identified by comparing their absence in fluorescence images to their presence in DIC images (Fig. 4 A). Rupture propensity is defined as the percentage of cells that undergo rupturing and it increases on increasing the strength of the hypo-osmotic shock. Rupture propensity also increases when the temperature is decreased from 37 °C to <10 °C or 25 °C or increased to 42 °C (Fig. 4 A, *right*). ATP depletion, too, increases rupture propensity, but only to 5-10% (Fig. 4 A, *right*). We find that RBCs (Fig. S3 A), in general, have a much higher rupture propensity than HeLa cells. In addition to calculating rupture propensity, we also follow the Calcein-AM loaded cells (HeLa and RBCs) after hypo-osmotic shock and find that rupturing events lead to a sudden loss in internal mean intensity (Figs. 4 B and S3 B). Ratio maps (Fig. 4 C) between consecutive images show that the rupturing is marked by fluorescence loss from the whole cell and by a simultaneous and sudden increase of fluorescence in the surrounding medium that is often asymmetric (Figs. 4C and S3 C). This indicates to a loss of intensity is due to a single-point rupture and is also seen in RBCs (Fig. S3 C, *left*). Fitting the temporal intensity profile (Figs. 4 D and S3 C, *right*) with exponential decay functions yields a time-constant which is used to estimate the rupture diameter based on a simple model that assumes fluorescence loss from the lesion by pure diffusion.

**Figure 4:**
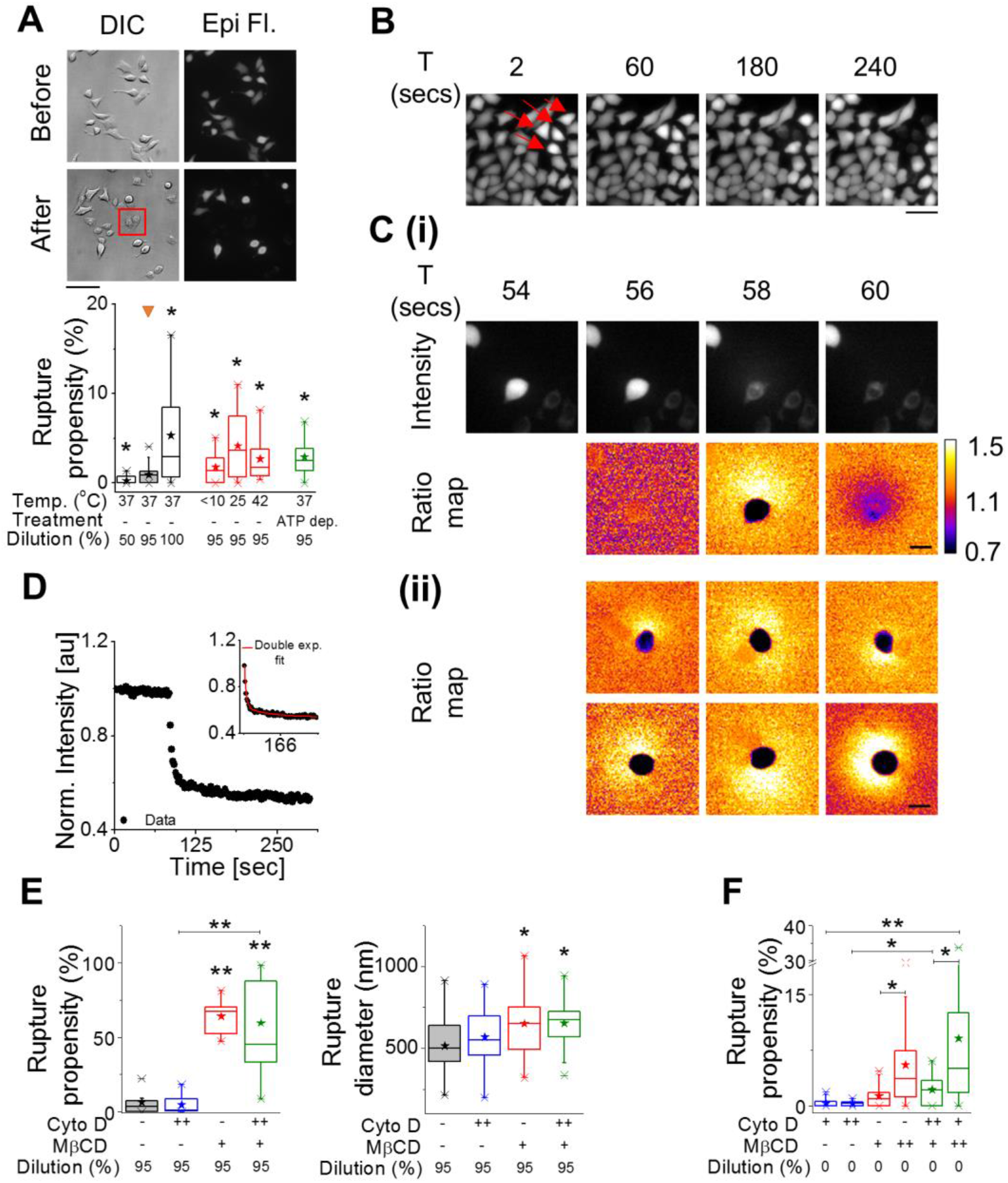
Membrane rupture induced by hypo-osmotic/iso-osmotic medium in the presence of MβCD. A) Top: DIC and epifluorescent images of Calcein AM labelled HeLa cells before and after the administration of a 95% hypo-osmotic shock (scale bar, 100 µm). Among others, the cells in the box are representatives of membrane rupture. Bottom: Box plot of rupture propensity of cells due to change in hypoosmotic stress, change in temperature and ATP depletion (N = 3 experiments each). B) Top: Time lapse images of Calcein AM loaded HeLa cells undergoing rupture (arrowheads in red) (scale bar, 50 µm). C) (i) Intensity and ratio 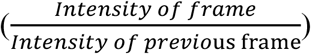 map of a rupturing cell followed in time to show single point rupture. (ii) Representative ratio maps of six different cells showing asymmetric spread of fluorescence after rupture (Scale bar, 30 µm). D) A time profile of normalized mean intensity of a cell; inset shows the double exponential fit to the profile. E) Box plots of rupture propensity (left) and rupture diameter (right) of 95% hypo-osmotic shock administered HeLa cells under control, Cyto D (2 hr post treatment), MβCD (50 mins post treatment) and Cyto D + MβCD conditions (N = 3 experiments each). F) Box plots of rupture propensity of cells under Cyto D, MβCD and dual drug treatments (N = at least 3 experiments each). ‘+’ denotes 60 mins treatment (for Cyto D), 50 mins (for MβcD), ‘++’ denotes 120 mins treatment. * p value < 0.05, ** p value < 0.001, Mann-Whitney U test. See Table S1 for statistics.

MβCD treatment results in an enhanced rupture propensity and an increased rupture diameter on hypo-osmotic shock (Fig. 4 E). As reported earlier ^15^, we too see that RBCs and a small percentage of HeLa cells rupture in isotonic media when treated with MβCD (Figs. 4 F and S3 E). We also find that increasing MβCD concentration enhances rupture propensity in RBCs while rupture diameter matches that with hypo-osmotically shocked control cells (Fig. S3 D, E).

Next, to understand the role of cytoskeleton in the measured effect, we perform experiments in which cells are first treated with Cyto D before MβCD treatment and then a hypo-osmotic shock is administered. We find that Cyto D, on its own does not alter the rupture propensity or rupture diameter on hypo-osmotic shock (Fig. 4 E). Similar values of rupture propensity and rupture diameter in MβCD treated cells as well as Cyto D + MβCD cells on hypo-osmotic shock show that the effect of MβCD is not abrogated by Cyto D. This indicates that the cytoskeleton is not essential for the effect of MβCD on membrane integrity. This effect is also seen in MβCD pre-treated cells (Fig. 4 F) and is found to be more pronounced in RBCs (Fig. S3 E and Movie S4).

Thus, cholesterol depletion by MβCD alters membrane mechanics by increasing the membrane tension, reducing the line and lysis tension which together enhance the rupturing propensity of the membrane with or without external stress. These effects are not mediated through the enhanced membrane-cytoskeleton interactions.

## DISCUSSION

In this paper, we use interference based membrane fluctuation maps to visualize and quantify the effect of MβCD mediated cholesterol depletion on cell membranes and their integrity. We report reduced temporal fluctuations and enhanced spatial flattening of the membrane on cholesterol depletion. Our study further focuses on the mechanics of the membrane – which, in contrast to earlier reports clearly shows that the membrane tension increases on cholesterol depletion. We also find that the interaction of the membrane with its underlying actin cytoskeleton increases, as reported previously ^7^.

Mapping fluctuations within single cells enables us to correlate the enhanced spatial heterogeneity in fluctuations with that in membrane tension. While fluctuations, in general, need not necessarily correlate with membrane tension, we show that the increased tension on MβCD treatment has a major contribution to the observed reduction in fluctuations. Even though the strength of the fluctuations is reduced, flattening out membrane undulations (increased correlation lengths) and reducing local non-uniformity (within 2.16 × 2.16 µm^2^ regions) do not abrogate signatures of active fluctuations in cholesterol depleted cells. But, when temporal fluctuation-amplitudes between different pairs of such regions located at different parts of the cell are compared, MβCD treated cells have more ‘dissimilar’ pairs than control cells. This implies that, in these cells, fluctuations can differ between membrane patches located at length scales larger than 2.16 µm. We further show that these regions are also differently tensed – the variability in tension being bigger than between ‘similar’ regions. Though the amplification of the long-range heterogeneity in fluctuations by MβCD treatment is robust in HeLa, it is not so in other cell lines (CHO or C2C12). However, the reduction in temporal fluctuations and its short-range variability, the increase in tension and its variability as well as the flattening of spatial undulations on cholesterol depletion are consistent in all the cell lines studied. These observations, together, clarify the effect of MβCD mediated cholesterol depletion on membrane mechanics and its spatial variability.

The ability to resist membrane rupture on hypo-osmotic shock is expected to be lost on cholesterol depletion ^14^ and is also seen in this study. However, our work addresses the role of membrane mechanics by quantifying the effect of MβCD on rupture propensity and diameter with and without hypo-osmotic shock. In cells with (HeLa) and without caveolae (RBCs), we find that MβCD aids ruptures, even in absence of external perturbation, implying that it destabilizes the membrane mechanically. The measured rupture diameter (r) corresponds to ratio of the membrane’s line to surface tension (γ/Σ) during lysis. Estimations show that the energy ^20,28^ required to open the pore 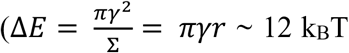 (for RBCs), 200 k_B_T (for HeLa), assuming a lower limit γ ∼ 1 pN and using observed radii of rupture, 15 nm and 250 nm, for RBC and HeLa cells respectively) is too high and contrasts the observed probability of rupture (expected (exp(-ΔE/ kBT)): 6x 10^−6^, observed: 0.092, for RBCs without hypo-osmotic shock and in the presence of MβCD). This implies that the ruptures could be induced by local defects in the membrane where there is a substantial reduction in line and lysis tension. While the exact values of tension cannot be extracted, we evaluate the relative changes in rupture diameter and propensity (Supporting Discussion) and argue that line ^20,45^ and lysis tension are reduced by MβCD in both RBCs and HeLa cells. It is possible that increased long-range spatial heterogeneity in fluctuations and membrane tension of MβCD treated cells reflect the existence of defects and that the increase in basal membrane tension by cholesterol depletion takes the system closer to the lowered lysis tension, hence aiding rupturing.

In conclusion, this work shows that under cholesterol depletion by MβCD, cells not only have altered fluctuations reflecting a flattening of spatial undulations but also shows a clear increase in membrane tension along with an increased long-range heterogeneity in fluctuations. Enhanced rupture rates and rupture diameter due to MβCD show that the membrane is also made vulnerable to rupture by a lowering of lysis and line tension.

## ACKNOWLEDGEMENTS

This work was supported by the Wellcome Trust/DBT India Alliance Fellowship (grant number IA/I/13/1/500885) awarded to Bidisha Sinha. We thank Rajesh Kumble Nayak for the code for calculating the PSD, Jayasri Das Sarma for HeLa cells, Rupak Datta for CHO cells and Kaushik Sengupta (Saha Institute of Nuclear Physics, Kolkata) for the C2C12 cells.

## AUTHOR CONTRIBUTIONS

Conceptualization, B.S.; Methodology, B.S., A.B., T.S., S.D., P.K.; Software, B.S., A.B.; Validation, A.B., B.S.; Investigation, A.B., P.K., S.D., T.S.; Formal Analysis, A.B., P.K.; Data Curation: A. B.; Writing – Original Draft, B.S., A.B.; Writing – Review & Editing, T.S., P.K.; Visualization, A.B.; Supervision, B.S.; Funding Acquisition, B.S.

## DECLARATION

The authors declare no competing financial interest.

